# Unipolar polysaccharide-mediated attachment of the N_2_O-reducing bacterium *Bradyrhizobium ottawaense* SG09 to plant roots

**DOI:** 10.1101/2025.06.05.658015

**Authors:** Yudai Takeguchi, Ryota Shibuya, Momoi Kondo, Eriko Betsuyaku, Manabu Itakura, Kiwamu Minamisawa, Masayuki Sugawara, Shigeyuki Betsuyaku

**Author notes:** **Corresponding authors’ contact information** Shigeyuki Betsuyaku, Masayuki Sugawara.

## Abstract

Agricultural soils are an important source of nitrous oxide (N_2_O), which has greenhouse and ozone-depleting effects. *Bradyrhizobium ottawaense* SG09 is a nitrogen-fixing rhizobium with high N_2_O-reducing activity. Rhizobia form symbiotic nodules in leguminous plants. The initial physical attachment of bacteria to plant roots is a critical step in the establishment of symbiotic interactions. In this study, we performed microscopic analysis using DsRed-expressing *B. ottawaense* SG09. We revealed that *B. ottawaense* SG09 attached to both the root surface and root hairs via single cellular poles. This polar attachment was observed not only to the symbiotic host soybean, but also to non-leguminous plants, such as *Arabidopsis*, rice, corn, and wheat. We identified and analyzed the *unipolar polysaccharide* (*upp*) gene cluster, which is proposed to be involved in polar attachment of rhizobia, in the genome of *B. ottawaense* SG09. We established an *Arabidopsis*-based interaction assay and demonstrated that *uppC* and *uppE* play a critical role in attachment to both the root surface and root hairs.

## INTRODUCTION

Nitrous oxide (N_2_O) is a long-lived greenhouse gas and ozone-depleting substance (Prather et al., 2015). It is released from natural and anthropogenic sources, with more than half of the latter originating from agricultural soils (Tian et al., 2020). *Bradyrhizobium ottawaense* SG09 is a nitrogen-fixing rhizobacterium with high N_2_O-reducing activity due to its high expression level of the *nosZ* gene, which encodes nitrous oxide reductase (Wasai-Hara et al., 2020, Wasai-Hara et al., 2023). The application of this bacterium is expected to effectively mitigate agricultural N_2_O emissions. However, attempts to infect legumes with specific rhizobial strains in agricultural fields often fail. This problem, which is due to competition between inoculants and indigenous rhizobia, highlights the importance of revealing the molecular basis of legume-rhizobia interactions (Thies et al., 1991).

To establish beneficial symbiosis with legumes, rhizobia form symbiotic structures called nodules where they fix nitrogen. This nodulation process is well studied in *Medicago truncatula* and *Lotus japonicus*. Specifically, flavonoid compounds secreted by plant roots activate the rhizobial NodD transcriptional activator, which induces production of Nod factors (Grundy et al., 2023). When host plant cells perceive Nod factors, infection threads form in root hairs, rhizobia enter these threads, and nodulation occurs (Gage. 2004). Prior to all these processes, the attachment of rhizobia to the host root surface is an important initial step. One of the most pronounced features of this process is the attachment of rhizobia to the cell surface of host roots via single cellular poles, as exemplified by several studies in *Agrobacterium tumefaciens* (Matthysse, 2014). Bacteria utilize various adhesive molecules to attach to surfaces (Knights et al., 2021). Among them, the polar adhesin unipolar polysaccharide (UPP), which is widely conserved in *Alphaproteobacteria*, is necessary for polar adhesion to both biotic and abiotic surfaces (Tomlinson and Fuqua 2008, Li et al., 2012, Fritts et al., 2017). Components of the UPP biosynthesis pathway are separately encoded by the core *upp* gene cluster, called *uppABCDEF*, and the others in the other part of the genomes. A similar structure called holdfast, which is conserved in *Caulobacterales*, is well studied in *Caulobacter crescentus* (Berne et al., 2015). Production of holdfast requires proteins encoded by the *hfs* gene cluster, called *hfsEFGHCBAD* (Smith et al., 2003, Toh et al., 2008). These two adhesins, UPP and holdfast, are distinct but share some similarities. For example, *uppC* and *uppD* show high sequence similarity to *hfsD* and *hfsE*, respectively (Fritts et al., 2017). Conservation of the *upp* gene cluster in plant-associated *Alphaproteobacteria* such as *A. tumefaciens* and *Bradyrhizobium japonicum* suggests that UPP is important for plant-microbe interactions (Xu et al., 2012, Xu et al., 2013, Fritts et al., 2017).

Several studies have demonstrated that UPP is required for unipolar binding of rhizobia to plant lectins *in vitro*. The plant symbiont *Rhizobium leguminosarum* is reported to utilize a UPP-type adhesin composed of glucomannan for attachment to pea and vetch roots, and this function is required for competitive nodule infection of pea roots (Laus et al., 2006; Williams et al., 2008, Knights et al., 2021). However, the role of the core *upp* gene cluster in *B. ottawaense* SG09 in the association with plants remains to be elucidated. In this study, we investigated the initial attachment of *B. ottawaense* SG09 to plant roots. We successfully visualized *B. ottawaense* SG09 using DsRed and revealed that this bacterium attached to the surface of root epidermal cells including root hairs via single cellular poles. This polar adhesion was observed not only in the interaction with the symbiotic host plant soybean, but also with non-leguminous plants, such as *Arabidopsis*, rice, corn, and wheat. We identified genes encoding the core *upp* gene cluster in the *B. ottawaense* SG09 genome and quantitatively evaluated their roles in the initial attachment to plant roots using deletion mutants of *uppC* and *uppE*.

## MATERIALS AND METHODS

### Bacterial strains and growth conditions

*B. ottawaense* strains were grown aerobically at 30°C in HM salt medium (Cole and Elkan, 1973) supplemented with 0.1% arabinose and 0.025% (w/v) yeast extract. *Escherichia coli* strains were grown at 37°C in Luria–Bertani medium. The following antibiotics were added appropriately: 300 mg L^−1^ neomycin (Nm) or 50 mg L^−1^ polymyxin B (Px) for *B. ottawaense* SG09 and 50 mg L^−1^ kanamycin for *E. coli*. DsRed-expressing *B. ottawaense* SG09 (SG09-DsRed) was generated using the pUT-based mini-Tn5 vector pBjGroEL4::DsRed2 carrying the DsRed2 sequence under the control of the BjGroEL4 constitutive promoter (Hayashi et al., 2014). All SG09-DsRed strains were grown in TY medium (5 g/L tryptone, 3 g/L yeast extract, and 1.3 g/L CaCl_2_・2H_2_O) containing 100 μg/mL spectinomycin at 28°C for 4–7 days.

### Construction of *uppE* and *uppC* deletion mutants

A mobilizable plasmid for generating the in-frame *uppE* deletion mutant (SG09DΔ*uppE*) was constructed as follows: a 0.9-kb fragment containing the 5’ flanking region of *uppE* (SG09_60820) and a fragment containing its 3′ flanking region were amplified by PCR from genomic DNA of *B. ottawaense* SG09 using the primer sets uppE_mutF1/uppE_mutR1 and uppE_mutF2/uppE_mutR2, respectively. The two PCR products were fused by overlap extension PCR, and the fused fragment was cloned into the *Sma*I site of pK18*mobsacB* (Schäfer et al., 1994) using an In-Fusion^®^ HD Cloning Kit (Takara Bio Inc., Kusatsu, Japan). The resulting plasmid (pMS195) was transferred from *E. coli* S17-1λpir to SG09-DsRed by biparental mating. An Nm/Px-resistant and sucrose-sensitive transconjugant was selected for single-crossover insertion of the plasmid into the chromosome. Cells were grown in HM liquid medium and spread on HM agar medium containing Px and 10% (w/v) sucrose to rescreen for sucrose-resistant colonies. The selected clones were further screened for Nm sensitivity. Double-crossover events and an in-frame deletion of *uppE* were confirmed by PCR using the primers uppE_F3/uppE_R3, and the nucleotide sequence of *uppE* in the mutants was determined by Eurofins Genomics (Tokyo, Japan) based on the Sanger method.

To generate the in-frame *uppC* deletion mutant (SG09DΔ*uppC*), a mobilizable plasmid was constructed as follows: a 0.7-kb fragment containing the 5’ flanking region of *uppC* (SG09_08120) and a fragment containing its 3′ flanking region were amplified by PCR from genomic DNA of *B. ottawaense* SG09 using the primer sets uppC_mutF1/uppC_mutR1 and uppC_mutF2/uppC_mutR2, respectively. The two PCR products were fused by overlap extension PCR, and the fused fragment was cloned into the *Sma*I site of pK18*mobsacB* as described above. The resulting plasmid (pMS196) was transferred from *E. coli* S17-1λpir to SG09-DsRed by biparental mating. The double-crossover mutants were selected as described above and confirmed by PCR using the primers uppC_F3/uppC_R3 and Sanger sequencing. The oligonucleotide primers used in this study are listed in Table S1.

### Plant growth conditions

Seeds of *Glycine max* cv. Enrei, *Arabidopsis thaliana* ecotype Col-0, *Oryza sativa* cv. Nipponbare, *Triticum aestivum* cv. Satonosora, and sweet corn (*Zea mays*) Wakuwaku Corn 82 Hybrid (Kaneko Seeds, Gunma, Japan) were used for microscopic analysis. Sterilized *Arabidopsis* seeds were plated and grown on 1/2 MS media (M0404; Sigma-Aldrich, St. Louis, Missouri, USA) containing 0.3% phytagel (P8169; Sigma-Aldrich, St. Louis, Missouri, USA) at 23°C with 24 h of light for the indicated number of days. The other seeds were sterilized and grown on 1.5% agar plates at 25°C for the indicated number of days.

### Confocal microscopy of bacterial interactions with various plant roots

Three- to five-day-old seedings were inoculated. Bacterial cells grown on TY media were resuspended in sterile water and adjusted to an optical density at 600 nm (OD_600_) of 0.003 for inoculation of soybean (4-day-old), rice (4-day-old), *Arabidopsis* (5-day-old), and wheat (3-day-old) and an OD_600_ of 0.03 for inoculation of sweet corn (3-day-old). The seedlings were incubated with 3 mL of the bacterial suspension at the indicated concentrations in 35-mm cell culture dishes with tissue culture-treated surfaces (3000-035; IWAKI, Shizuoka, Japan). The roots of overnight-treated seedlings were excised, washed once with sterile water, and mounted between the glass surface of a glass-bottom dish (3910-035, IWAKI) and a 1/2 MS phytagel block. Fluorescence confocal and reflection imaging were performed using an LSM980 Airyscan 2 confocal microscope equipped with an LD LCI Plan-Apochromat 40×/1.2 Imm Corr DIC M27 objective and ZEN software (Carl Zeiss Microscopy, Oberkochen, Germany).

### Quantification of bacterial attachment to the root surface

Five-day-old Col-0 seedings were inoculated. Bacterial cells were prepared as described above. Six seedlings were submerged in 10 mL of the bacterial suspension (OD_600_=0.003) in an Aznol Petri Dish JP, φ90×15 mm (AS ONE, Osaka Japan) and incubated in this suspension at 23°C for 24 h in the dark. Prior to microscopic analysis, seedlings were washed twice with sterile water. The root surface area located 5 mm from the first root hair that appeared from the shoot side was selected for image acquisition using a BX53 fluorescence microscope equipped with a UPLFLN 40× objective and cellSens Standard software (Evident, Tokyo Japan) (Fig. S1). A circle with a radius of 50 μm centered on the root in the microscopic image was selected as the count area, but lateral roots and lateral root primordia were excluded, using Fiji (a distribution of ImageJ, version 2.14.0/1.54p) (Fig. S1A). Experiments were performed with six biological replicates and repeated three times individually.

### Quantification of bacterial attachment to root hairs

Four-day-old Col-0 seedings were inoculated. Bacterial inoculation was performed as described above. After inoculation, the seedlings were incubated at 23°C for 3 h in the dark. Prior to microscopic analysis, plants were washed twice with sterile water. An area of the root centered 2.5 mm from the first root hair from the root tip side was selected for image acquisition using a BX53 fluorescence microscope equipped with a UPLFLN 10× objective and cellSens Standard software (Evident) (Fig. S1B). A square measuring 1 × 1 mm^2^ centered on the root within the microscopic images was selected for counting using Fiji (a distribution of ImageJ, version 2.14.0/1.54g) (Fig. S1B). The number of bacteria per root hair was counted, and root hairs were classified into three groups based on the bacterial count: 0 (class I), 1–5 (class II), and 6 and more (class III). Experiments were performed with 3–6 biological replicates and repeated three times individually.

## RESULTS

### *B. ottawaense* SG09 exhibits unipolar binding to the root surface

To gain insights into the initial physical interaction between *B. ottawaense* SG09, which has high N_2_O-reducing activity, and plant root tissues, we focused on soybean, a potential agricultural target for practical application of this bacterium. Rhizobial attachment to legume roots is affected by environmental factors such as soil pH (Knights et al., 2021). To simplify the comparison, we developed a simple incubation assay using aseptically germinating soybean seedlings together with *B. ottawaense* SG09 carrying chromosomally integrated DsRed2 under the control of a constitutive promoter (SG09-DsRed) in sterile water. Confocal laser scanning microscopy and confocal reflection microscopy (CRM) were employed to visualize DsRed fluorescence from SG09-DsRed cells and non-fluorescently labeled plant tissue structures, respectively, allowing reconstruction of the three-dimensional (3D) structure of bacterial attachment to plant tissues (Paddock, 2002; Kiyokawa et al., 2017). After incubation for 24 h, the roots were washed and analyzed. Confocal laser scanning microscopy and CRM revealed that multiple SG09-DsRed cells adhered to the soybean root epidermal surface via single poles, resembling well-known characteristics of rhizobia (Fig. 1A and Supplementary Movie 1) (Fritts et al., 2017). An orthogonal projection of the reconstituted 3D structure clearly visualized unipolar attachment of individual SG09-DsRed cells to the plant cell surface (Fig. 1B). Next, we examined if this unipolar adhesion to the plant cellular surface is specific to soybean, which *B. ottawaense* SG09 establishes a symbiotic relationship with through nodulation (Wasai-Hara et al., 2020). When similarly incubated with the roots of rice, wheat, sweet corn, and even *Arabidopsis*, SG09-DsRed cells attached to all the roots examined via single cellular poles (Fig. 2, Fig. S2, Supplementary Movies 2 and 3). Such unipolar attachment was not only observed to root epidermal surfaces but also to root hairs (an example of rice root hairs is shown in Fig. 2D and E, Supplementary Movie 4). These results showed that *B. ottawaense* SG09 adhered to the cellular surfaces of various plant species without any specific preference.

**Fig. 1.**
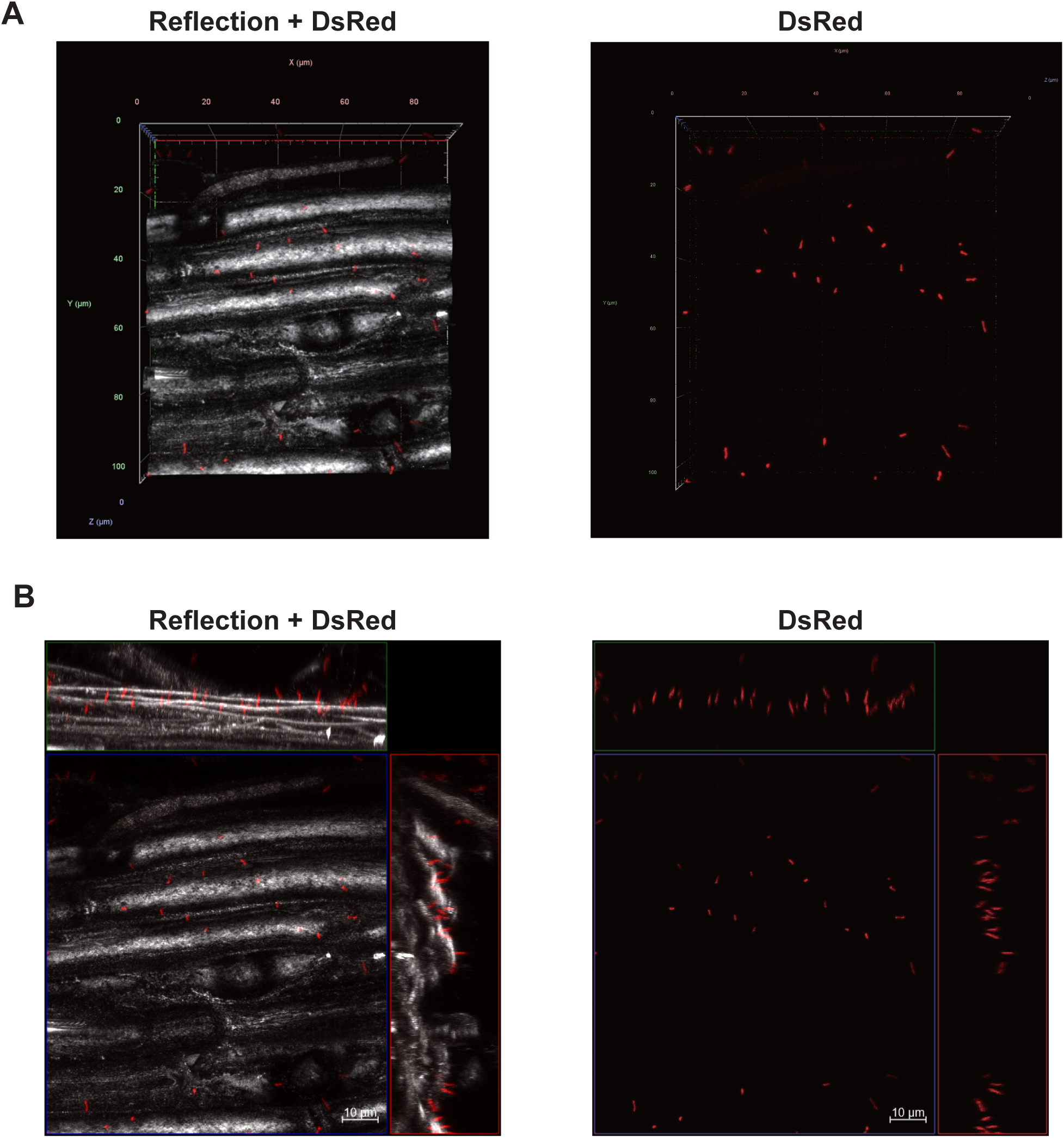
Confocal fluorescence imaging of the roots of soybean seedlings incubated with SG09-DsRed. (A) Representative reconstituted 3D images of SG09-DsRed (red) binding to soybean roots visualized by CRM (gray). A merged image (left) and the corresponding DsRed image (right) are shown. (B) Orthogonal projection images of the reconstituted 3D image in (A) represented by maximal projection. A merged image (left) and the corresponding DsRed image (right) are shown. Scale bar, 10 µm.

**Fig. 2.**
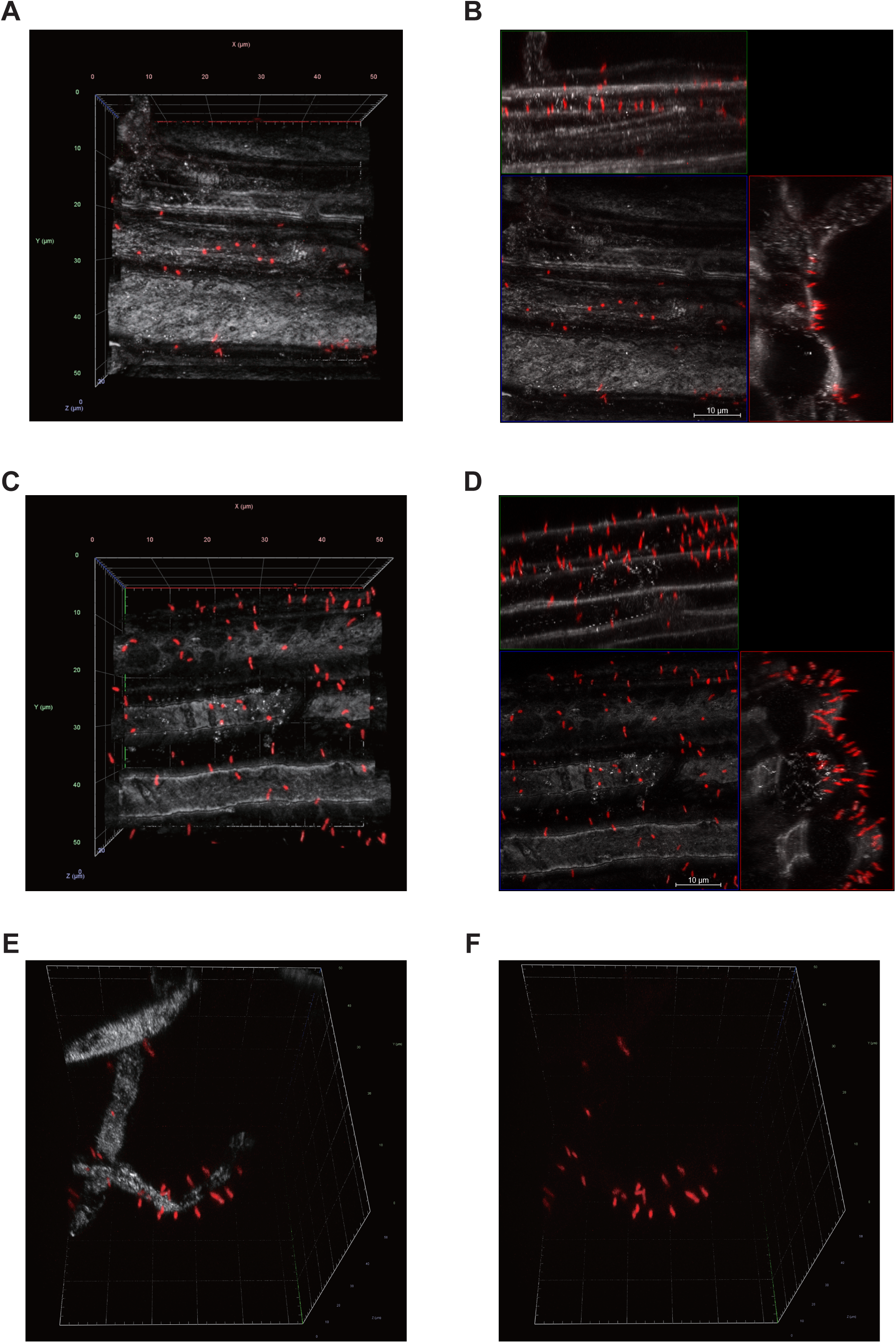
Confocal fluorescence imaging of the roots of rice and *Arabidopsis* seedlings incubated with SG09-DsRed. (A) A representative reconstituted 3D image of SG09-DsRed (red) binding to rice roots visualized by CRM (gray). (B) An orthogonal projection image of the reconstituted 3D image in (A) represented by maximal projection. (C) A representative reconstituted 3D image of SG09-DsRed (red) binding to *Arabidopsis* roots visualized by CRM (gray). (D) An orthogonal projection image of the reconstituted 3D image in (C) represented by maximal projection. (E) Representative reconstituted 3D images of SG09-DsRed (red) binding to rice root hairs visualized by CRM (gray). (F)A DsRed image of the 3D images shown in (E). Scale bars, 10 µm.

### The core *upp* gene cluster is conserved in the *B. ottawaense* SG09 genome

UPP was previously reported to be required for bacterial polar attachment; therefore, we investigated whether *B. ottawaense* SG09 possesses homologous genes to *uppABCDEF* of *R. palustris* CGA009 (Fritts et al., 2017; Onyeziri, et al., 2022). BLASTP analysis identified homologous genes in the *B. ottawaense* SG09 genome for each of the *R. palustris* CGA009 genes, and these homologs were consequently designated *uppA*, *uppB*, *uppC*, *uppD*, *uppE*, and *uppF* in *B. ottawaense* SG09 (Fig. 3A, Table S2). The core *upp* gene cluster in the *B. ottawaense* SG09 genome consists of *uppABDEF*, while *uppC* is located separately, showing synteny with that of *B. japonicum* USDA 110 (Fritts et al., 2017). *uppC* and *uppE* in *R. palustris* and *A. tumefaciens* play a crucial role in production of UPP; therefore, we generated SG09-DsRed in-frame deletion mutants of *uppC* and *uppE*, called SG09DΔ*uppC* and SG09DΔ*uppE* (Fig. 3B and C) (Fritts et al., 2017; Onyeziri, et al., 2022).

**Fig. 3.**
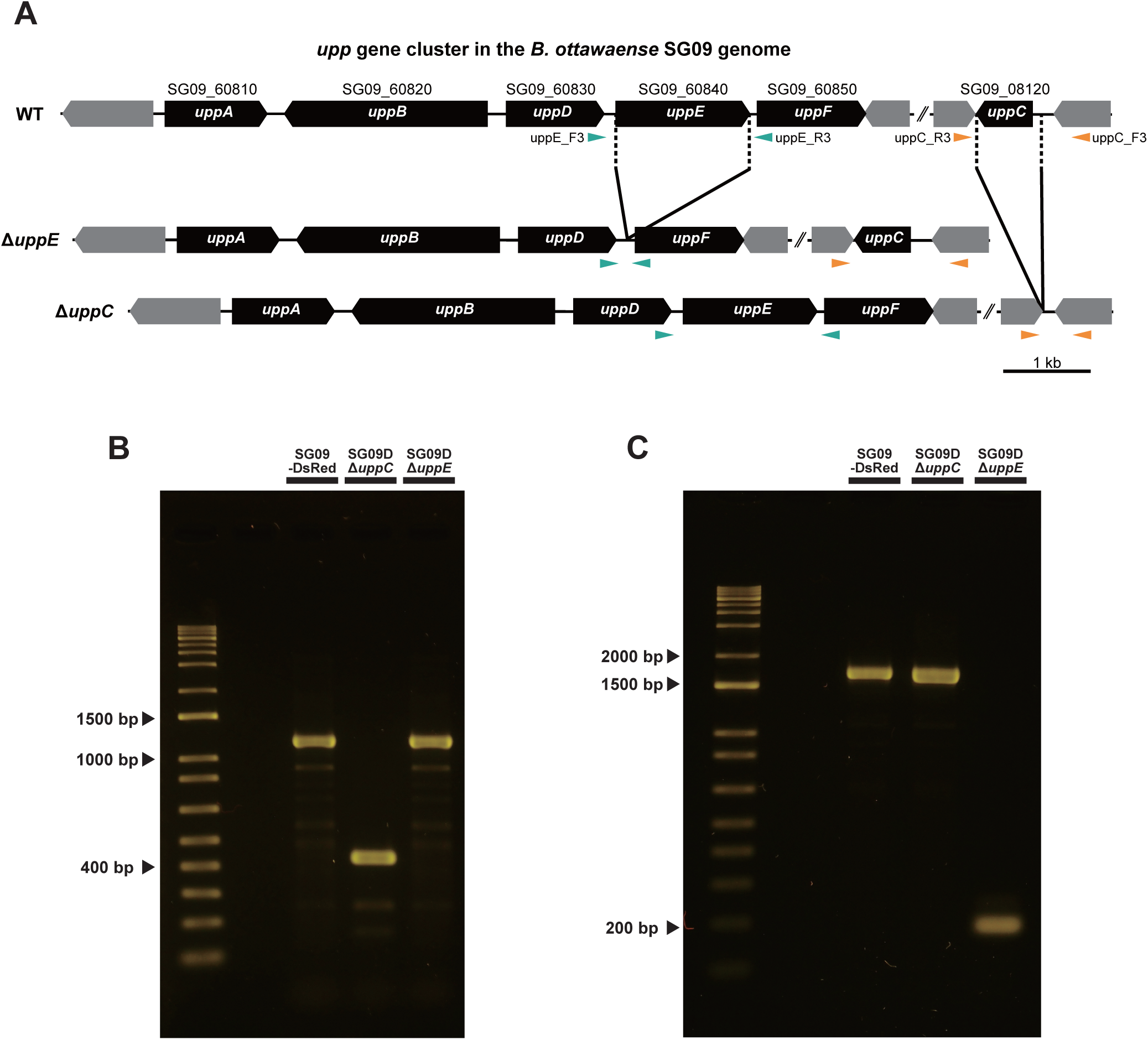
The core *upp* gene cluster in the *B. ottawaense* SG09 genome and the mutants generated in this study. (A) Schematic representations of the *upp* gene cluster in the genomes of wild-type *B. ottawaense* SG09 and the respective mutants. (B, C) PCR validation of SG09DΔ*uppC* and SG09DΔ*uppE* loci using the primer pairs uppC_F3/uppC_R3 for *uppC* (B) and uppE_F3/uppE_R3 for *uppE* (C). PCR products were separated on 2% agarose gels.

### *uppC* and *uppE* are required for attachment of *B. ottawaense* SG09 to plant surfaces

Functional analysis of bacterial mutants in terms of their polar attachment is often performed using *in vitro* assays such as the biofilm formation and lectin-binding assays. The use of cultured soybean cells and a specific antibody against a *B. japonicum* strain enabled a more direct and quantitative binding assay with plant cells, but it is costly and labor-intensive to raise specific antibodies and maintain cell cultures (Ho et al., 1988). Considering that SG09-DsRed exhibits unipolar attachment to roots of a small plant, *Arabidopsis*, in addition to soybean, we microscopically investigated the roles of *uppC* and *uppE* using *Arabidopsis* ecotype Col-0 as a host in a small-scale, direct, and quantitative attachment assay. We designed two assays to characterize SG09DΔ*uppC* and SG09DΔ*uppE*, along with SG09-DsRed, in terms of epidermal and root hair attachment (see Materials and Methods). Although the number of SG09-DsRed that attached to the root surface varied greatly among root samples, significantly fewer SG09DΔ*uppC* and SG09DΔ*uppE* than SG09-DsRed attached (Fig. 4). Additionally, significantly fewer SG09DΔ*uppC* than SG09DΔ*uppE* attached to the root surface (Fig. 4).

**Fig. 4.**
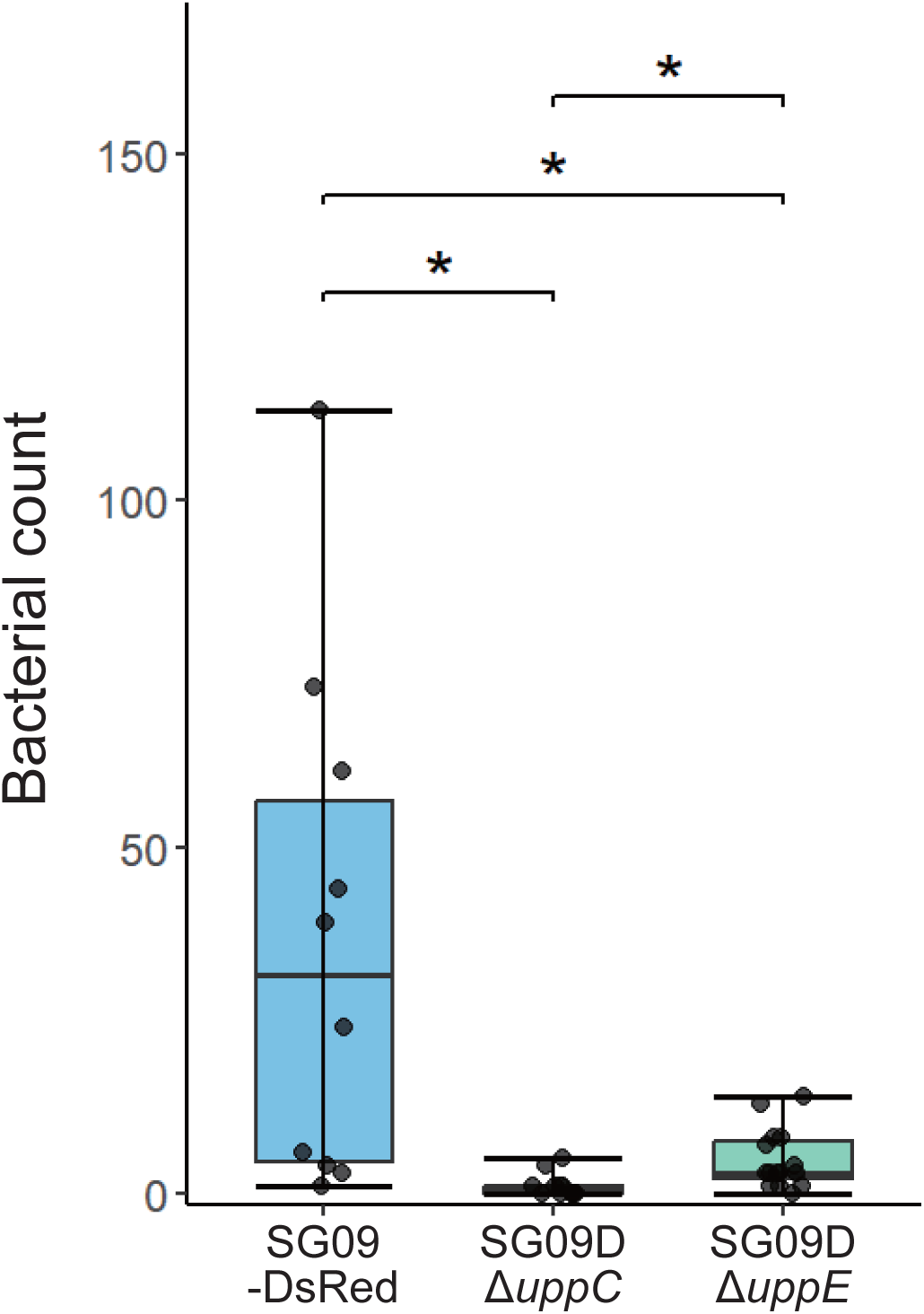
Quantification of bacterial attachment to the root surface. Five-day-old Col-0 seedlings were incubated with bacterial suspensions (OD_600_=0.003). After incubation for 24 h at 23°C in the dark, plants were washed twice with sterile water. The number of epidermally attached bacterial cells present within a circle with a radius of 50 μm set 5 mm away from the first root hair observed from the shoot side was counted. *n*=10 for SG09-DsRed, *n*=13 for SG09DΔ*uppC*, and *n*=15 for SG09DΔ*uppE*. \**p* < 0.05. Statistical analysis was performed by the *t*-test using R (version 4.4.2).

Next, we focused on root hair attachment. There was great variation in the number of bacteria that attached to root hairs; therefore, root hair attachment was analyzed by classifying the observed phenotypes into three categories based on the number of attached bacterial cells: 0 (class I), 1–5 (class II), and 6 and more (class III). Compared with SG09-DsRed, both mutants exhibited a significant increase in the number of class I root hairs (Fig. 5A) and a significant reduction in the number of class III root hair (Fig. 5C). In terms of the numbers of class I and class III root hairs, SG09DΔ*uppC* had a significantly more severe phenotype than SG09DΔ*uppE* (Fig. 5A and C). The number of class II root hairs did not significantly differ among the tested strains, suggesting that this class reflects accidental attachment of bacterial cells to root hairs (Fig. 5B). Taken together, our quantitative microscopic analysis reveals that UppC and UppE play a direct and essential role in attachment of *B. ottawaense* SG09 cells to the root epidermal surface and root hairs.

**Fig. 5.**
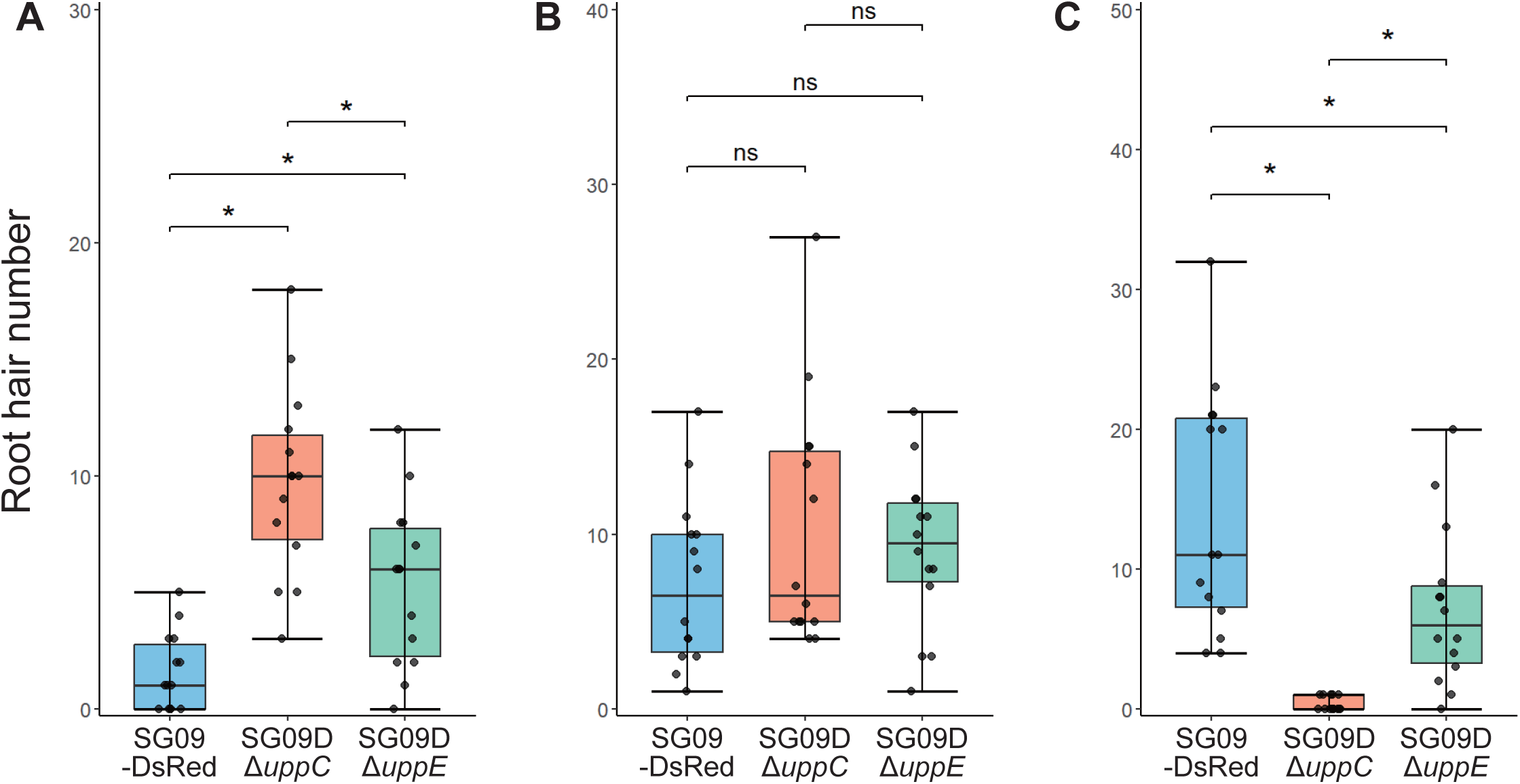
Quantification of bacterial attachment to root hairs. Four-day-old Col-0 seedlings were incubated with bacterial suspensions (OD_600_=0.3). After incubation for 3 h at 23°C in the dark, plants were washed twice with sterile water. A one mm square centered 2.5 mm from the first root hair from the root tip side was used for bacterial counting. The number of bacterial cells attached to each root hair was counted and then root hairs are classified into three groups based on the number of attached bacteria: 0 (A, class I), 1–5 (B, class II), and 6 and more (C, class III). *n*=14. \**p* < 0.05; ns, not significant. Statistical analysis was performed by the *t*-test using R (version 4.4.2).

## DISCUSSION

In this study, we demonstrated that *B. ottawaense* SG09 attached to both the root surface and root hairs via single cellular poles. This attachment was observed not only to soybean, a symbiotic leguminous host for this bacterium, but also to non-leguminous host plants, such as *Arabidopsis*, rice, corn, and wheat. We identified that the *upp* gene cluster is conserved in the *B. ottawaense* SG09 genome. Disruption of *uppC* and *uppE* in *B. ottawaense* SG09 significantly reduced attachment of this bacterium to both the root surface and root hairs in *Arabidopsis*. Our assay using *Arabidopsis* provides a high-throughput approach *in planta*. In addition, it enables us to use several genetic tools, *e.g.,* various well-known mutants of plant lectins and immune system components, to molecularly elucidate the interactions of *B. ottawaense* SG09 with plants.

There is a marked phenotypic difference between SG09DΔ*uppC* and SG09DΔ*uppE*, suggesting that UppC plays a crucial role in attachment to both the root surface and root hairs. *uppC* and *uppE* encode the Wza-type outer membrane polysaccharide secretin and WbaP-initiating glycosyltransferase, respectively, in *A. tumefaciens* (Onyeziri et al., 2022). In *A. tumefaciens*, *uppE* is necessary for UPP production under inorganic phosphate-rich conditions, while *uppE* and its paralog *Atu0102* act redundantly under inorganic phosphate-limited conditions (Xu et al., 2012). Thus, other genes might function redundantly with *uppE* in the *B. ottawaense* SG09 genome. Our findings suggest that UPP is important for attachment of *B. ottawaense* SG09 to biotic surfaces. Further information is essential to elucidate the function of UPP in *B. ottawaense* SG09, *e.g.,* the biochemical composition that governs lectin specificity (Onyeziri et al., 2022). However, it is difficult to determine the biochemical composition of UPP because of its insolubility and low level of synthesis by bacterial cells (Berne et al., 2015, Thompson et al., 2018).

*B. ottawaense* SG09 is gaining attention because of its high N_2_O-reducing activity (Wasai-Hara et al., 2023). Further optimization is required to exploit *B. ottawaense* SG09 as an N_2_O-reducing agent in agricultural fields, including overcoming the competitive effects of indigenous bacteria (Thies et al., 1991, Wasai-Hara et al., 2023). The initial process of bacterial attachment to the root surface could be an important target for inoculated bacteria to successfully dominate during nodulation. In this study, we established a small-scale, direct, and quantitative *in planta* assay using *Arabidopsis* as a host, which could broaden our knowledge of the mechanisms underlying rhizobial attachment. Further research may lead to engineering of rhizobial adhesins that improve the probability of attachment and the development of methods that overcome indigenous bacteria.

## Acknowledgements

This work was supported by project JPNP18016, subsidized by the New Energy and Industrial Technology Development Organization (NEDO).

## REFERENCES

Cole, M. A. & Elkan, G. H. Transmissible resistance to penicillin G, neomycin, and chloramphenicol in *Rhizobium japonicum*. Antimicrob Agents Chemother 4, 248–253 (1973).

Berne, C., Ducret, A., Hardy, G. G., & Brun, Y. V. (2015). Adhesins Involved in Attachment to Abiotic Surfaces by Gram-Negative Bacteria. Microbiology spectrum, 3(4), 10.1128.

Fritts, R. K., LaSarre, B., Stoner, A. M., Posto, A. L., & McKinlay, J. B. (2017). A Rhizobiales-Specific Unipolar Polysaccharide Adhesin Contributes to *Rhodopseudomonas palustris* Biofilm Formation across Diverse Photoheterotrophic Conditions. Applied and environmental microbiology, 83(4), e03035–16.

Gage D. J. (2004). Infection and invasion of roots by symbiotic, nitrogen-fixing rhizobia during nodulation of temperate legumes. Microbiology and molecular biology reviews : MMBR, 68(2), 280–300.

Grundy, E. B., Gresshoff, P. M., Su, H., & Ferguson, B. J. (2023). Legumes Regulate Symbiosis with Rhizobia via Their Innate Immune System. International journal of molecular sciences, 24(3), 2800.

Hayashi, M., Shiro, S., Kanamori, H., Mori-Hosokawa, S., Sasaki-Yamagata, H., Sayama, T., et al. (2014) A Thaumatin-Like Protein, Rj4, Controls Nodule Symbiotic Specificity in Soybean. Plant Cell Physiol. 55: 1679–1689.

Ho, S.C., Ye, W.Z., Schindler, M., and Wang, J.L. (1988) Quantitative assay for binding of Bradyrhizobium japonicum to cultured soybean cells. J Bacteriol. 170: 3882–3890.

Kiyokawa, T., Usuba, R., Obana, N., Yokokawa, M., Toyofuku, M., Suzuki, H., et al. (2017) A Versatile and Rapidly Deployable Device to Enable Spatiotemporal Observations of the Sessile Microbes and Environmental Surfaces. Microbes Environment. 32: ME16161.

Knights, H. E., Jorrin, B., Haskett, T. L., & Poole, P. S. (2021). Deciphering bacterial mechanisms of root colonization. Environmental microbiology reports, 13(4), 428–444.

Laus, M. C., Logman, T. J., Lamers, G. E., Van Brussel, A. A., Carlson, R. W., & Kijne, J. W. (2006). A novel polar surface polysaccharide from *Rhizobium leguminosarum* binds host plant lectin. Molecular microbiology, 59(6), 1704–1713.

Li, G., Brown, P. J., Tang, J. X., Xu, J., Quardokus, E. M., Fuqua, C., & Brun, Y. V. (2012). Surface contact stimulates the just-in-time deployment of bacterial adhesins. Molecular microbiology, 83(1), 41–51.

Onyeziri, M. C., Hardy, G. G., Natarajan, R., Xu, J., Reynolds, I. P., Kim, J., Merritt, P. M., Danhorn, T., Hibbing, M. E., Weisberg, A. J., Chang, J. H., & Fuqua, C. (2022). Dual adhesive unipolar polysaccharides synthesized by overlapping biosynthetic pathways in *Agrobacterium tumefaciens*. Molecular microbiology, 117(5), 1023–1047.

Paddock, S. (2002) Confocal reflection microscopy: the “other” confocal mode. Biotechniques. 32: 274, 276–8.

Prather, M. J., Hsu, J., DeLuca, N. M., Jackman, C. H., Oman, L. D., Douglass, A. R., Fleming, E. L., Strahan, S. E., Steenrod, S. D., Søvde, O. A., Isaksen, I. S., Froidevaux, L., & Funke, B. (2015). Measuring and modeling the lifetime of nitrous oxide including its variability. Journal of geophysical research. Atmospheres : JGR, 120(11), 5693–5705.

Schäfer, A. et al. Small mobilizable multi-purpose cloning vectors derived from the *Escherichia coli* plasmids pK18 and pK19: selection of defined deletions in the chromosome of *Corynebacterium glutamicum*. Gene 145, 69–73 (1994).

Smith, C. S., Hinz, A., Bodenmiller, D., Larson, D. E., & Brun, Y. V. (2003). Identification of genes required for synthesis of the adhesive holdfast in *Caulobacter crescentus*. Journal of bacteriology, 185(4), 1432–1442.

Thies, J. E., Singleton, P. W., & Bohlool, B. B. (1991). Influence of the size of indigenous rhizobial populations on establishment and symbiotic performance of introduced rhizobia on field-grown legumes. Applied and environmental microbiology, 57(1), 19–28.

Tian, H., Xu, R., Canadell, J. G., Thompson, R. L., Winiwarter, W., Suntharalingam, P., Davidson, E. A., Ciais, P., Jackson, R. B., Janssens-Maenhout, G., Prather, M. J., Regnier, P., Pan, N., Pan, S., Peters, G. P., Shi, H., Tubiello, F. N., Zaehle, S., Zhou, F., Arneth, A., … Yao, Y. (2020). A comprehensive quantification of global nitrous oxide sources and sinks. Nature, 586(7828), 248–256.

Toh, E., Kurtz, H. D., Jr, & Brun, Y. V. (2008). Characterization of the *Caulobacter crescentus* holdfast polysaccharide biosynthesis pathway reveals significant redundancy in the initiating glycosyltransferase and polymerase steps. Journal of bacteriology, 190(21), 7219–7231.

Tomlinson, A. D., & Fuqua, C. (2009). Mechanisms and regulation of polar surface attachment in *Agrobacterium tumefaciens*. Current opinion in microbiology, 12(6), 708–714.

Thompson, M. A., Onyeziri, M. C., & Fuqua, C. (2018). Function and Regulation of *Agrobacterium tumefaciens* Cell Surface Structures that Promote Attachment. Current topics in microbiology and immunology, 418, 143–184.

Wasai-Hara, S., Hara, S., Morikawa, T., Sugawara, M., Takami, H., Yoneda, J., Tokunaga, T., & Minamisawa, K. (2020). Diversity of *Bradyrhizobium* in Non-Leguminous Sorghum Plants: *B. ottawaense* Isolates Unique in Genes for N2O Reductase and Lack of the Type VI Secretion System. Microbes and environments, 35(1), ME19102.

Wasai-Hara, S., Itakura, M., Fernandes Siqueira, A., Takemoto, D., Sugawara, M., Mitsui, H., Sato, S., Inagaki, N., Yamazaki, T., Imaizumi-Anraku, H., Shimoda, Y., & Minamisawa, K. (2023). *Bradyrhizobium ottawaense* efficiently reduces nitrous oxide through high *nosZ* gene expression. Scientific reports, 13(1), 18862.

Williams, A., Wilkinson, A., Krehenbrink, M., Russo, D. M., Zorreguieta, A., & Downie, J. A. (2008). Glucomannan-mediated attachment of *Rhizobium leguminosarum* to pea root hairs is required for competitive nodule infection. Journal of bacteriology, 190(13), 4706–4715.

Xu, J., Kim, J., Danhorn, T., Merritt, P. M., & Fuqua, C. (2012). Phosphorus limitation increases attachment in *Agrobacterium tumefaciens* and reveals a conditional functional redundancy in adhesin biosynthesis. Research in microbiology, 163(9-10), 674–684.

Xu, J., Kim, J., Koestler, B.J., Choi, J., Waters, C.M., and Fuqua, C. (2013) Genetic analysis of Agrobacterium tumefaciens unipolar polysaccharide production reveals complex integrated control of the motile[to[sessile switch. Mol Microbiol. 89: 929–948.

