## Supplementary material for "Unipolar polysaccharide-mediated attachment of the N_2_O-reducing bacterium *Bradyrhizobium ottawaense* SG09 to plant roots": Fig. S

### Figure S1.

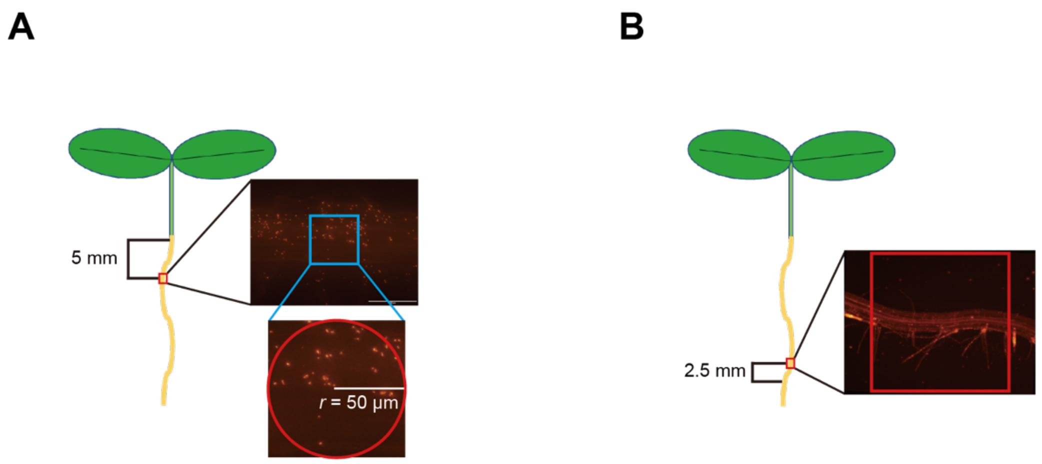


Figure S1. Schematic diagrams showing the measurement methods used in this study. (A) Root surface attachment analysis. A circle with a radius of 50 μm centered on the root (shown in red) located 5 mm from the first root hair in the microscopic image was used for bacterial counting. (B) Root hair attachment analysis. The number of bacterial cells per root hair in a one mm square centered on the root (shown in red) located 2.5 mm from the first root hair from the root tip side was used for bacterial counting.

**Figure S2.**

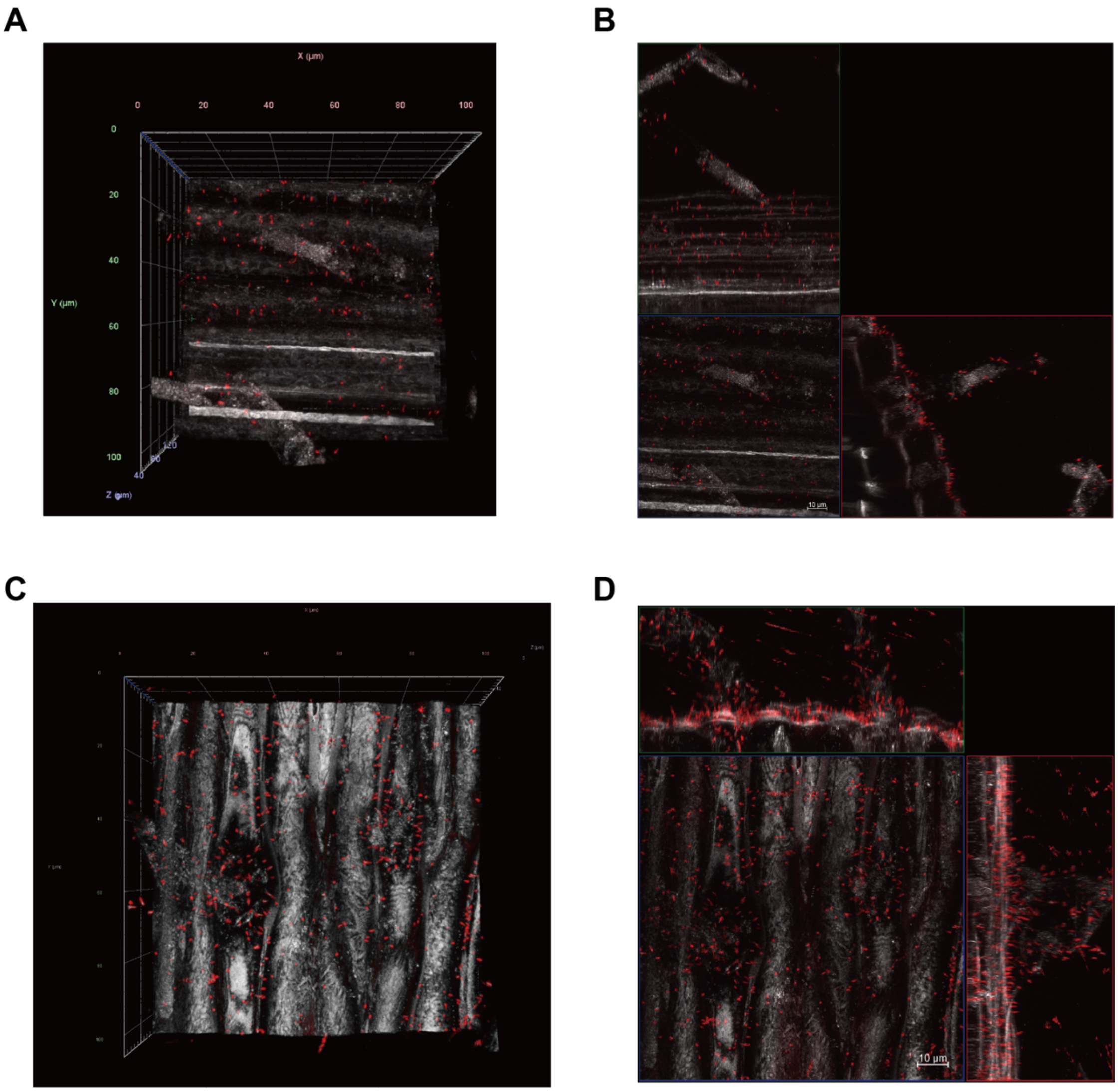


Figure S2. Confocal fluorescence imaging of the roots of wheat and sweet corn seedlings incubated with SG09-DsRed.

(A) A representative reconstituted 3D image of SG09-DsRed (red) binding to wheat roots visualized by CRM (gray). (B) An orthogonal projection image of the reconstituted 3D image in (A) represented by maximal projection. (C) A representative reconstituted 3D image of SG09-DsRed (red) binding to sweet corn roots visualized by CRM (gray). (D) An orthogonal projection image of the reconstituted 3D image in (C) represented by maximal projection. Scale bars, 10 µm.

Supplementary Movie 1. Reconstituted 3D rendering of SG09-DsRed (red) binding to soybean roots visualized by CRM (gray) shown in Fig. 1A.

Supplementary Movie 2. Reconstituted 3D rendering of SG09-DsRed (red) binding to rice roots visualized by CRM (gray) shown in Fig. 2A.

Supplementary Movie 3. Reconstituted 3D rendering of SG09-DsRed (red) binding to *Arabidopsis* roots visualized by CRM (gray) shown in Fig. 2C.

Supplementary Movie 4. Reconstituted 3D rendering of SG09-DsRed (red) binding to rice root hairs visualized by CRM (gray).

Table S1.

**Table S1. Primers used in this study**

| Primer name | Sequence (5' to 3') |
|---|---|
| uppC_mutF1 | TCGAGCTCGGTACCCCGACCACCCATTTCGTCAAC |
| uppC_mutR1 | TGGATTTCAGGCGGTGAGCGAGACGGCTTAAGTGATGC |
| uppC_mutF2 | GCATCACTTAAGCCGTCTCGCTCACCGCCTGAAATCCA |
| uppC_mutR2 | CTCTAGAGGATCCCCTTTGAAATAGGGCGCGTCTC |
| uppC_F3 | CCTATTCCGATGCGCAGAAG |
| uppC_R3 | CATCGACGCTAACAGGATCG |
| uppE_mutF1 | TCGAGCTCGGTACCCCGCTGCATTATCCCCTCAAC |
| uppE_mutR1 | CGGGGCATGACGGAAAACTCCACGTCCATTTCAAGCAC |
| uppE_mutF2 | GTGCTTGAAATGGACGTGGAGTTTTCCGTCATGCCCCG |
| uppE_mutR2 | CTCTAGAGGATCCCCGTGGAATCGAACGACAGCAG |
| uppE_F3 | TAACCATCCTTAAGCCCGCT |
| uppE_R3 | CTATTCGGACGCCAAACACC |

*The underlined sequences are homologous regions of pK18mobsacB.

Table S2.

**Table S2. BLASTP analysis of the orthologous genes of *uppABCDEF* in *B. ottawaense* SG09**

| ***Rhodopseudomonas palustris* CGA009** | ***Bradyrhizobium ottawaense* SG09** | | |
|---|---|---|---|
| | **Locus tag** | **Position** | **Identity(%)** |
| RPA2753_UppA | SG09_60810 | 6502658-6503839 | 230/390 (59%) |
| RPA2752_UppB | SG09_60820 | 6504021-6506357 | 484/699 (69%) |
| RPA2751_UppD | SG09_60830 | 6506563-6507696 | 262/377 (69%) |
| RPA2750_UppE | SG09_60840 | 6507817-6509364 | 401/519 (77%) |
| RPA4581_UppF | SG09_60850 | 6509436-6510692 | 227/404 (56%) |
| RPA4833_UppC | SG09_08120 | 888458-889111 | 143/217 (66%) |
